# Time is of the essence: A general framework for uncovering temporal structures of communities

**DOI:** 10.1101/2023.06.30.546877

**Authors:** Hannah Yin, Volker H. W. Rudolf

## Abstract

Ecological communities are inherently dynamic: species constantly turn over within years, months, weeks, or even days. These temporal shifts in community composition determine essential aspects of species interactions and how energy, nutrients, information, diseases, and perturbations "flow" through systems. Yet, our understanding of community structure has relied heavily on static analyses not designed to capture critical features of this dynamic temporal dimension of communities. Here, we propose a conceptual and methodological framework for quantifying and analyzing this temporal dimension. Conceptually, we split the temporal structure into two definitive features, sequence and duration, and review how they are linked to key concepts in ecology. We then outline how we can capture these definitive features using perspectives and tools from temporal graph theory. We demonstrate how we can easily integrate ongoing research on phenology into this framework and highlight what new opportunities arise from this approach to answer fundamental questions in community ecology. As climate change reshuffles ecological communities worldwide, quantifying the temporal organization of communities is imperative to resolve the fundamental processes that shape natural ecosystems and predict how these systems may change in the future.

## The temporal dimension of community structure

Communities are organized across two fundamental dimensions: time and space. Research on spatial patterns has increased rapidly in the past decades and transformed how we think about natural communities by emphasizing the importance of spatial organization (structure) of species (Kareiva 1994; Brown 1995; Leibold & Chase 2017; Tilman & Kareiva 2018). In comparison, we still know surprisingly little about how communities are organized in time, which has only recently gained renewed attention as a line of research (Wolkovich *et al*. 2014; Post 2019; Yang 2020). The ongoing and projected impacts of climate change on ecological communities, as well as calls for their short- and long-term mitigation and adaptation plans (IPCC 2023), create an urgency to develop frameworks explicitly focusing on the temporal dimension of community structure to fill this conceptual gap (Yang & Rudolf 2010; Wolkovich *et al*. 2014; Rudolf 2019; Takimoto & Sato 2020; Yang 2020; CaraDonna *et al*. 2021).

Ecologists have long recognized that the temporal patterns and shifts in community composition are a hallmark of any natural community (Elton 1927). Their importance manifests in several critical ecological concepts, like the temporal niche (Schoener 1974; Carothers & Jaksić 1984; Loreau 1989a; Jensen *et al*. 2019), succession (HA 1927; Connell & Slatyer 1977; Poorter *et al*. 2023), the window of opportunity (Yang *et al*. 2008; Yang & Cenzer 2020), and the seasonal timing of life history events (phenology) (Parmesan & Yohe 2003; Miller-Rushing *et al*. 2010; Cohen *et al*. 2018; Post 2019; Roslin *et al*. 2021). Yet, despite the widespread appreciation of the dynamic nature of community structure, community ecology has traditionally taken a largely static approach. For instance, the structure of ecological networks is typically assumed to be fixed and time-invariant such that only the abundance of species may change over time (Montoya *et al*. 2006; Ings *et al*. 2009; CaraDonna *et al*. 2021). When analyzing empirical patterns, studies of community structure predominantly use a single data "snapshot" in time or aggregate time points to calculate key biodiversity metrics, thereby potentially overlooking transient states and critical ecological processes present at the community level, such as seasonality (McMeans *et al*. 2015). Consequently, our current understanding of the temporal structure of natural communities remains limited concerning its variations across ecosystems, space, and time and the factors that drive such variations. Yet, answering these and many related questions about the temporal structure of communities is fundamental to our ability to understand and predict the dynamics of natural communities and how they will respond to environmental change (Wolkovich *et al*. 2014; Rudolf 2019; Yang 2020; CaraDonna *et al*. 2021).

Given that ecologists have long appreciated the dynamic structure of communities, what has prevented us from answering these questions already? In large part, progress has been hampered by the lack of a unified conceptual and methodological framework for analyzing the temporal patterns and the logistical challenges in obtaining the fine-scaled temporal data required, which can require a lot of resources or new technologies. To move toward filling these gaps, we propose a general framework to quantify and analyze temporal structure in natural communities. First, we define the concept of temporal community structure and discuss its potential role in both well-studied and understudied ecological concepts. We then briefly introduce temporal- ordered networks and suggest how and why these networks can theoretically capture the definitive features of temporal structure in natural communities by applying perspectives from temporal graph theory. We end by applying this framework to publicly available phenological data and highlight some promising research directions and outstanding questions that this approach helps address.

### What is the temporal structure of communities?

Just like "spatial structure" generally denotes how species are organized in space, "temporal structure" refers to the organization of species in time. However, unlike space, time is unidirectional: it always moves forward, never stops, and we can’t go back. This directionality means that events and processes from the past can affect events and processes that happen in the future, but not the other way around. Temporal structure is thus fundamentally about the *sequence* (chronological order) and *duration* of and between events (Wolkovich *et al*. 2014).

Take a piece of music, for example. Knowing all the notes played in a melody does not tell you how it sounds. What makes any melody unique is the sequence of notes (relative timing), how long each note is, and the pauses between them. Similarly, knowing what species exist in a community (aggregated over time) doesn’t reveal much about how the community looks at any given point in time or the "pace" or "rhythm" of key ecological processes and change. Instead, if we are to truly understand the ecological and evolutionary processes that shape the dynamics of natural communities and how communities will respond to environmental change, we need to study their temporal structure: we need to analyze the duration and sequence of when certain groups (e.g., species) are active and their intervening times (gaps).

### Why are the sequence and duration of events so important?

The critical aspects of temporal structure—sequence and duration of and between events— play essential roles in structuring species interactions and ecosystem processes. The sequence and duration of species’ phenophases determine which species co-occur and, thus, what interactions are possible. A predator cannot consume prey, and a bee cannot pollinate a flower if neither is present at the right stage at the right time(s). Understanding what interactions are possible through temporal co-occurrences can thus help predict and capture species interactions, even though co-occurrence does not necessarily imply a direct interaction between species (Thurman *et al*. 2019; Blanchet *et al*. 2020). For instance, the loss of temporal overlap of phenologies can explain the loss of plant-pollinator interactions over a century (Burkle *et al*. 2013). Similarly, species with longer phenophases tend to co-occur with more species and thus have more species interactions (Resasco *et al*. 2021) (**Fig. 1a**). However, in this case, interactions also typically occurred in sequence with only one or a few interactions at any given point in time. Thus, species that might have seemed like generalists were often “sequential specialists” (**Fig. 1b**). These examples highlight that the sequence and duration of species’ activities can help identify the possible interactions and reveal the forces that structured life histories and community composition.

**Figure 1.**
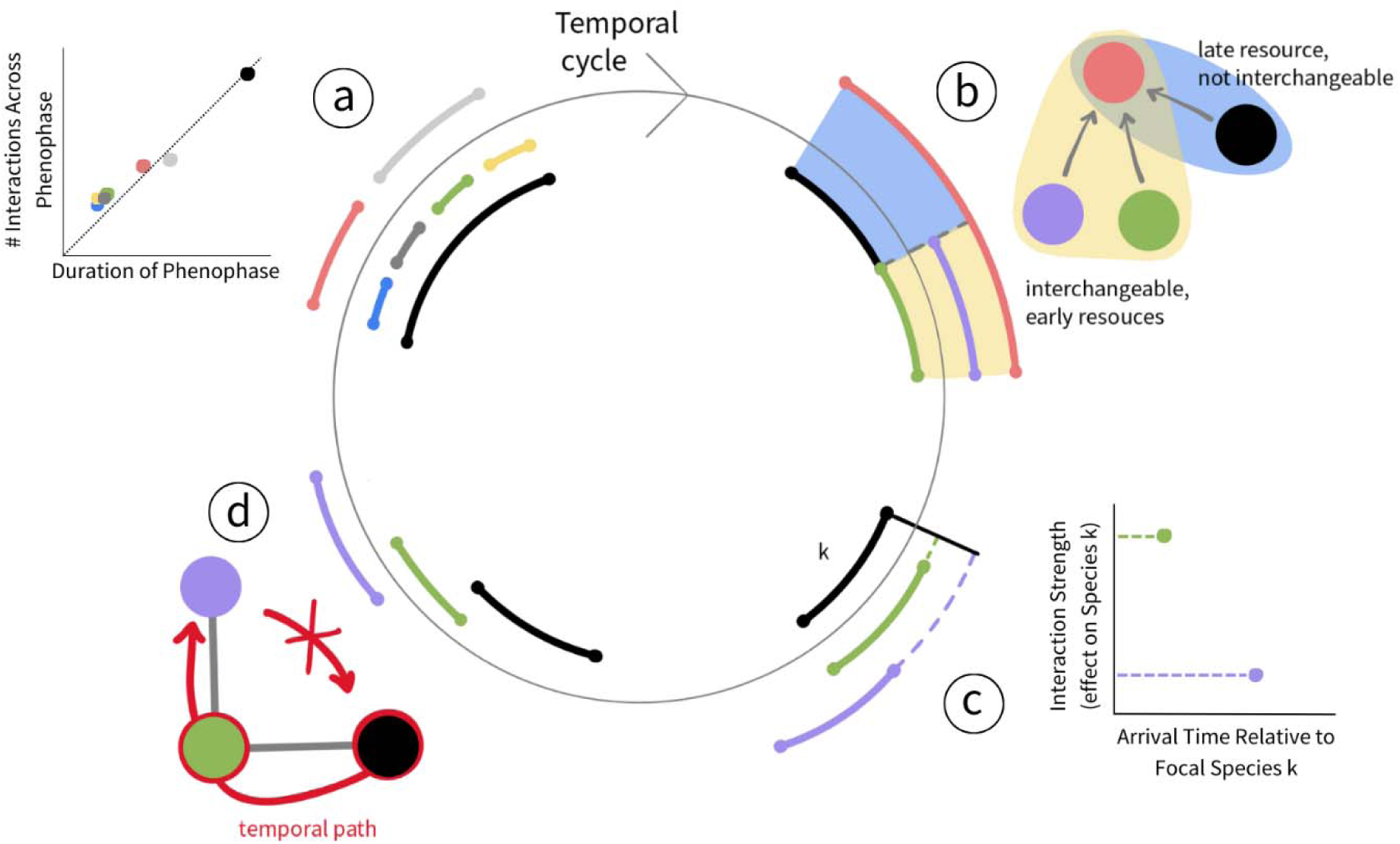
Temporal structure, as defined by the sequence and duration of and between events, influences many aspects of species interactions. Our four examples (a-d) may happen simultaneously or in different orders and were arranged arbitrarily along a cyclical rather than linear timeline to emphasize that many natural events reoccur periodically. Line segments indicate the presence of a species for a temporal (e.g., annual) cycle. Colors represent different species. **(a)** How long species are present (length of phenophases) typically correlates positively with the number of interactions per species (e.g., the black species has more interaction than the green or the grey species). **(b)** Consumers who appear as generalists in static networks (coral) often feed on resources that occur during different periods (purple, green, black) and thus are not interchangeable. **(c)** The sequence of species’ arrival times and duration between arrivals can determine the strength of species’ interactions. **(d)** The order of phenologies determines what direct and indirect interactions are possible in a community: early-arriving species (black) can affect later-arriving species, even after the early arrivers leave (e.g., via resource depletion, habitat modification, or other community members), but not vice versa.

The sequence of species arrival times and duration between arrivals can also determine the outcome and strength (i.e., per-capita effect) of species interactions (Rudolf 2019; Zou & Rudolf 2023) (**Fig. 1c**). For instance, reducing the overlap of flowering times can either increase or decrease the reproduction of plant species in a community via recruitment or competition for pollinators respectively (Mitchell *et al*. 2009; Faust & Iler 2022). Early arrivers often have a competitive advantage over species that arrive later, and this advantage typically depends on the difference in relative arrival times (Rasmussen *et al*. 2014; Rudolf 2018; Alexander & Levine 2019; Blackford *et al*. 2020). Similarly, predation rates decline with a delay in salamander and dragonfly predator arrival relative to its tadpole prey (Rudolf 2022b). Such timing effects (also called priority effects or historical contingencies) are well documented in animal, plant, and microbial systems and determine the dynamics and composition of natural communities (Fukami 2015; Zou & Rudolf 2023).

Furthermore, the sequence and spacing in which species are arranged along a temporal axis also determines the causal chain of possible events (Blonder *et al*. 2012). In the absence of temporal overlap, species present earlier in a community can still affect those that appear later but not vice versa (**Fig. 1d**). For instance, experiments in pond communities found that *Bufo* tadpoles that breed early in the season can negatively affect later arriving *Hyla* tadpoles, even though the two species never co-occurred in time (Wilbur & Alford 1985). Such "legacy effects" are common in natural communities and can arise through a range of different mechanisms (Fukami 2015), e.g., by altering the environment (Kostenko *et al*. 2012), consuming a competitor of a later arriving species, or by changing the community composition that later arriving species experience (Rudolf & Van Allen 2017). However, the potential to affect other community members naturally declines towards the end of a season; e.g., species that arrive at the end of a season tend to have fewer interactions (i.e., are more specialized) (Seifert *et al*. 2021). The species that arrive last can only affect a limited number of species with which they co-occur. The relative temporal position of species is, therefore, inherently linked to its potential impact on the community and how it can be affected by other community members.

At the community and ecosystem level, the duration and sequence of temporal co-occurrences limit the possible pathways through which diseases, energy, water, organic and inorganic elements, and perturbations cascade through the ecosystem. Perturbations (e.g., fire, drought, heatwave, chemical spill, or disease outbreak) that happen in the middle of a season can only affect species that are already present or arrive later in the season, while species that arrived and left before the perturbation will not be affected. Thus, explicitly accounting for the sequence and duration of events is imperative to capture the mechanisms that drive community dynamics and composition, ecosystem processes, and how they will respond to environmental change.

### What can we learn from the temporal structure of communities?

The presence of species along the temporal axis is not random but rather an emergent property that reflects its ecological and evolutionary responses to constantly changing environmental conditions and interactions. Therefore, the temporal positions of species can tell us a lot about the ecology of species and the processes that structure natural communities. For example, specific patterns of temporal spacing between species presences can suggest functional relationships of species, competition and coexistence mechanisms, or predict the invasion success of species (**Table 1**). Anomalies in temporal spacing may reveal when environmental conditions create windows of opportunities or constraints (Yang & Rudolf 2010; Yang 2020; Yang & Cenzer 2020) and indicate facilitative interactions or co-dependence (Dante *et al*. 2013). As mentioned earlier, temporal co-occurrence indicates, at minimum, what interactions are possible and is, therefore, a crucial driver of the structure of ecological networks (Yang *et al*. 2013; Morente-López *et al*. 2018; CaraDonna *et al*. 2021). Temporal structure also helps us to predict the potential pathways and even speed of transmission of energy, infectious diseases, and perturbations (Blonder 2015). These examples and studies highlight how temporal structure has contributed to our understanding of community processes.

**Table 1:** Examples of questions in community ecology linked to temporal patterns.

| Ecological concept | Temporal pattern used to infer concepts | Example references |
| --- | --- | --- |
| What are potential mechanisms of community assembly? | Reduced temporal variation & non-random temporal patterns (over time, e.g., years or habitats) were used to infer the strength of stochastic versus deterministic processes, historical contingencies, and priority effects. | (Purschke *et al.* 2013; Van Allen *et al.* 2017; Neale & Rudolf 2023) |
|  | Slope of STR (species time relationship) can be used to infer processes that determine biodiversity, like dispersal/migration & extinction rates, succession, and speciation, or to differentiate neutral from niche based processes. | (Rosenzweig 1995; McKinney & Frederick 1999; Adler 2004; White *et al.* 2006; Rivett *et al.* 2021) |
| What are potential coexistence mechanisms? | The sequence of species' arrivals and time between them determines the potential for coexistence (via correlated changes in fitness and niche differences). | (Godoy & Levine 2014; Alexander & Levine 2019; Rudolf 2019; Blackford *et al.* 2020) |
| Do species compete for and partition time? | Lower temporal overlap than expected from a null model was used to infer temporal niche partitioning and competition. | (Loreau 1989b; Koide *et al.* 2007; Jensen *et al.* 2019; Schamp & Jensen 2019) |
|  | Altered phenologies after species removal suggested reduced competition, via temporal niche separation. | (Dumandan *et al.* 2023) |
| What is the functional relationship between species? | Temporal covariances of occurrences helped infer species interactions. | (Steele *et al.* 2011; Ushio *et al.* 2018; Deutschmann *et al.* 2023) |
|  | Temporal clustering was used to infer facilitation/positive indirect interactions between species. | (Dante *et al.* 2013) |
|  | Phenological tracking can indicate strong inter-dependency of species (e.g., predator-prey, plant-pollinator). | (Visser & Gienapp 2019) |
|  | Competitive dominance depends on the relative time between and order of species arrivals. | (Rudolf 2018; Rudolf & McCrory 2018; Faust & Iler 2022) |
|  | Phenological timing determines predator-prey and host-parasitoid interactions and community structure. | (Alford 1989; Pardikes *et al.* 2022; Rudolf 2022a) |
| What factors determine network structure? | Loss of temporal overlap results in loss of interactions. | (Burkle *et al.* 2013) |
|  | Phenological attributes determine network robustness and | (Ramos-Jiliberto *et al.* 2018; Guzman |
|  | persistence of species. | *et al.* 2021) |
|  | Phenology drives modularity. | (Morente-López *et al.* 2018) |
| What determines invasion success? | Invasion success/growth rates dependent on the relative timing of arrival (ex: phenology of germination or flowering). | (Wolkovich & Cleland 2011; Godoy & Levine 2014; Zettlemoyer *et al.* 2019) |
| How do traits determine specialization? | Species with narrow time windows (short presence) are more likely to specialize on temporary resources and certain environmental conditions (e.g., seasonal specialist versus generalist). | (Humphries *et al.* 2017) |
| What determines transmissions of pathogens, nutrients, energy? | Temporal structure predicts the possible pathways and speed of transmission through a community (e.g., faster with higher average overlap, less time between arrival). | (Blonder *et al.* 2012) |
| How can we predict the effects of perturbations? | The effects of perturbations on species and communities depend on the specific timing of perturbation and temporal organization of species. | (Crawley 2004; Craine *et al.* 2012; Shepherd *et al.* 2012; Li & Pennings 2017) |
| How can we identify windows of opportunity? | Temporal clustering and gaps in species presences can indicate favorable (and unfavorable) environmental conditions. | (Martinez Falcon et al 2010) |

Nevertheless, we still have significant gaps in our current understanding of temporal community structure. Ecology still lacks a general framework to predict what temporal structure should look like and how it is shaped by and interacts with various processes. For instance, how do changes in season length, productivity, biodiversity, alter the duration and sequence of species presences and how species partition time? What patterns of the temporal structure of communities are typically robust and shared across most ecosystems and which tend to vary? We can start to answer these and many related questions by analyzing and comparing temporal structure across taxa, systems, environments, space, and timescales to help us identify and understand many biological processes that shape natural communities. However, because the past can shape the future, a mechanistic and comprehensive understanding of temporal patterns must account for the sequence of events to capture causality. Importantly, this exploration encourages the development of more well-rounded and nuanced frameworks for the roles both space and time play in communities and more testable and accurate predictions of how they will respond to environmental change (Wolkovich *et al*. 2014; Yang 2020).

Finally, studying the temporal structure of communities also provides new quantitative approaches to detect phenological shifts (e.g., due to climate change) and put them in a community context. Current metrics typically do not capture concurrent changes in the temporal structure of entire communities. Yet, as we will explain in more detail below, quantifying changes in the temporal structure of communities can increase the power to detect phenological shifts and help us learn more about their ecological consequences.

## The need for a general framework to quantify temporal community structure

Because various studies and related ecological concepts use the sequence and duration of (or between) events to understand community dynamics (**Table 1**), we believe in the merits of establishing a general quantitative framework designed to capture the critical features of temporal community structure. Current approaches to analyzing the temporal structure of communities fall mainly into a few research areas: phenology, species turnover (including succession), and interaction networks. Each approach has strengths and limitations in capturing the complexity and essential features of the temporal structure of communities.

To date, phenology studies have predominantly focused on metrics for the onset (timing) or duration of phenological events at the single species level (Forchhammer *et al*. 1998; CaraDonna *et al*. 2014; Cohen *et al*. 2018), pairwise correlations of these events (Ovaskainen *et al*. 2013; Thackeray *et al*. 2016), and the temporal overlap of interacting pairs of species (Visser & Both 2005; CaraDonna *et al*. 2014; Theobald *et al*. 2017; Carter *et al*. 2018; Renner & Zohner 2018; Visser & Gienapp 2019). While these and other studies have been fundamental in documenting the temporal coordination and variability of interactions over time, many aspects of the temporal community structure only emerge from the community context. For instance, the dynamic structure of communities could show periods of high activity separated by periods of inactivity. We can only identify such bursts of activity by analyzing the distribution of "gaps" between phenologies of all species. Furthermore, distributions of multiple species’ phenologies and gaps between them could characterize distinct temporal modules, e.g., subsets of interacting species that always co-occur or a set of species in a module to consistently appear in the same chronological order. Of course,while co-occurrence does not necessarily mean the realization of their direct interactions, co-occurrence is a necessary condition. Such modules may thus help predict the presence/absence and strength of interactions, symbiotic relationships, temporal guilds, temporal niche partitioning, or identify temporal windows of opportunity and species that exploit similar resources or abiotic conditions (**Table 1**). Similarly, the ecological consequences of phenological shifts may only become fully apparent in a community context (e.g., when shifts create temporal partitioning, **Fig. 2**). In short, the temporal features characteristic of species’ phenologies may suggest important community dynamics, but identifying these temporal features requires a holistic approach that moves beyond analyses of single-species or pairwise metrics that we traditionally use in phenological studies.

**Figure 2.**
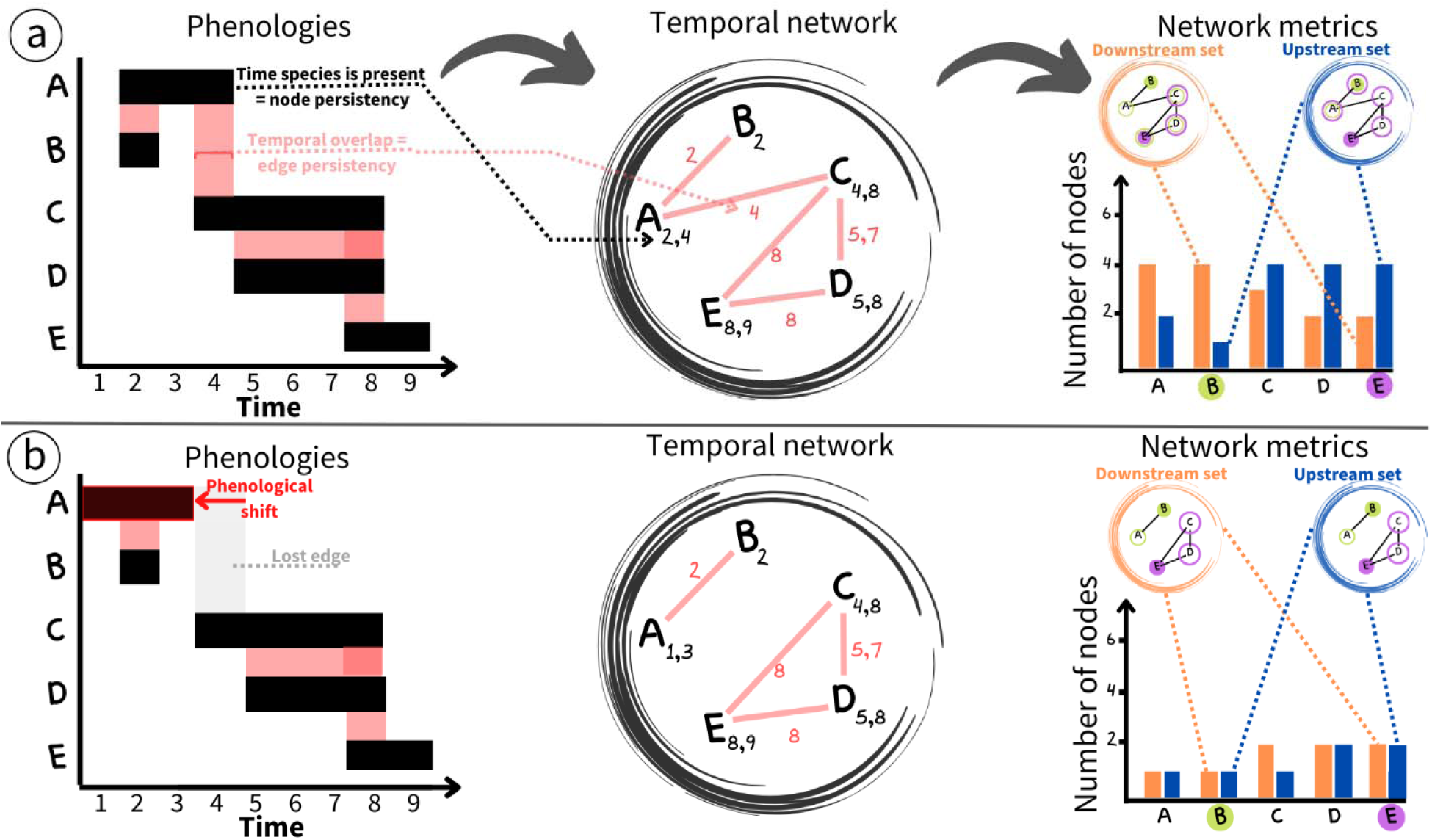
Steps to build, analyze, and compare time-ordered networks using phenology data. The start and end date of a phenophase quantifies when species are present and active in a community (black bars in left column). We can then calculate the species’ temporal overlap/co-occurrence (red bars). From a network perspective, the length of the phenophase (black bar) and overlap (red bar) indicate the persistency of nodes and links, respectively. We use this phenology data to build time-ordered networks of communities (middle column), where nodes are species with corresponding time stamps that indicate when a species is present. Time stamps can be a single time point, a time interval with start and end date, or multiple points or intervals if species re-enter a community. The temporal co-occurrence of species gives links that connect nodes (species) with their corresponding time stamps (red bars). Once these time-ordered networks are constructed, we can calculate a range of time-ordered network metrics that explicitly account for the temporal sequence of events (see **Table 2**). For instance, we can find the "downstream set" and "upstream set" for each species (bar graphs on the right) to determine how species influence each other given the temporal sequence of species occurrences/phenologies (see **Table 2** for complete definitions). Circular inserts show a graphical example of how those two metrics are calculated for focal species B and E, with a solid circle indicating a focal species and an open circle indicating a corresponding (by color) source or set of influence species. These metrics are sensitive to the relative temporal position of a species and the sequence of temporal overlap. They provide new insights into the basic temporal structure of communities and can also be used to gain a deeper understanding of the functional role of species. For instance, species that appear earlier often have a more extensive downstream set of nodes but a smaller upstream set than species that appear late (e.g., B vs E in the top right panel). However, this can change when temporal gaps (no species present) are present, which create "subsets" of networks (top vs bottom bar graph). (**a**) versus (**b**) shows how a phenological shift in a single species can lead to dramatic changes in network structure and metrics, especially when this results in the loss of a link that connects temporally separated groups of species (e.g., a shift of species A in top scenario (**a**) versus scenario (**b**)), highlighting the utility of this approach for comparing temporal structure of communities across space and time and quantifying phenological shifts in a community context.

**Table 2:**
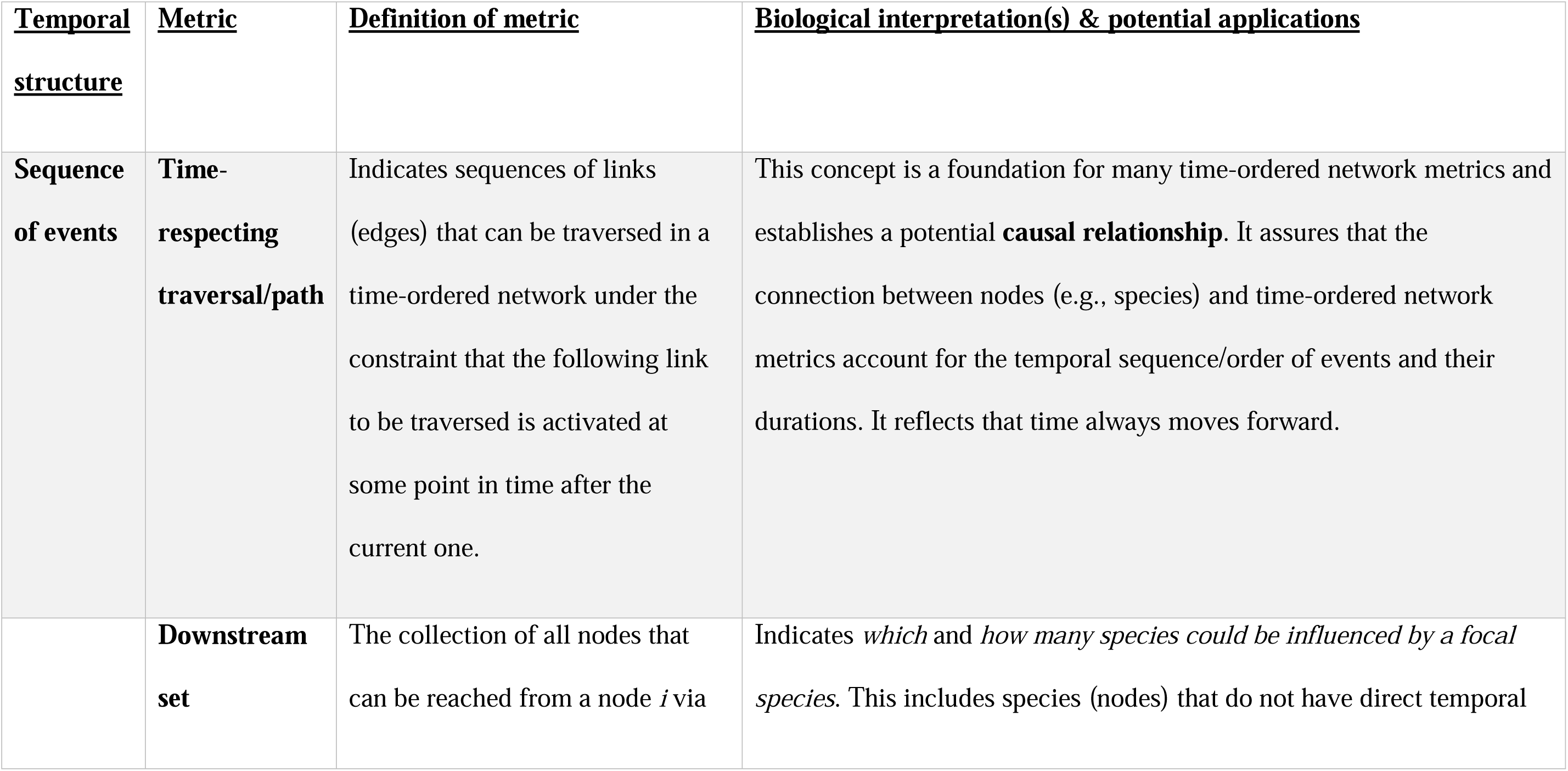
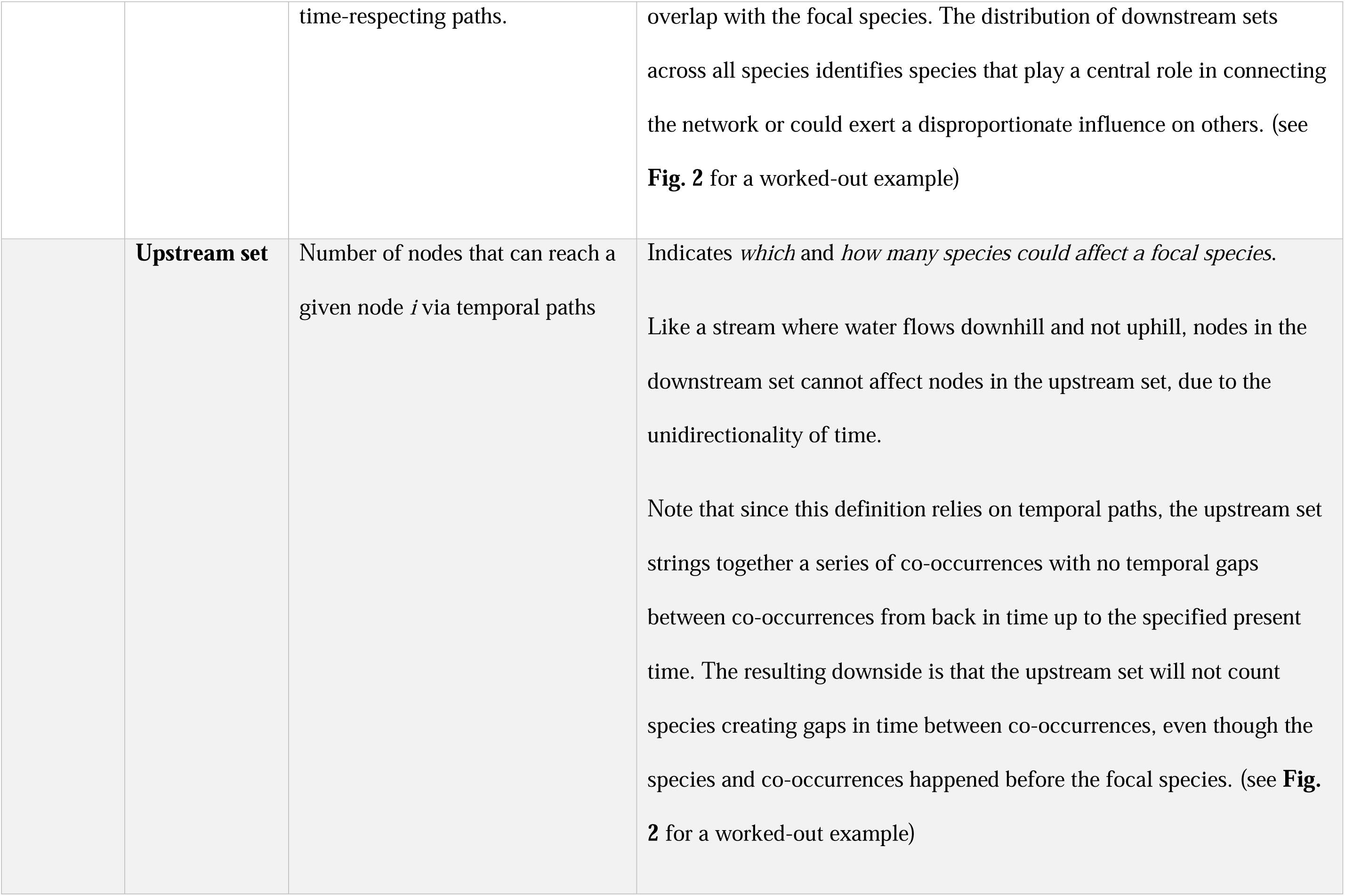
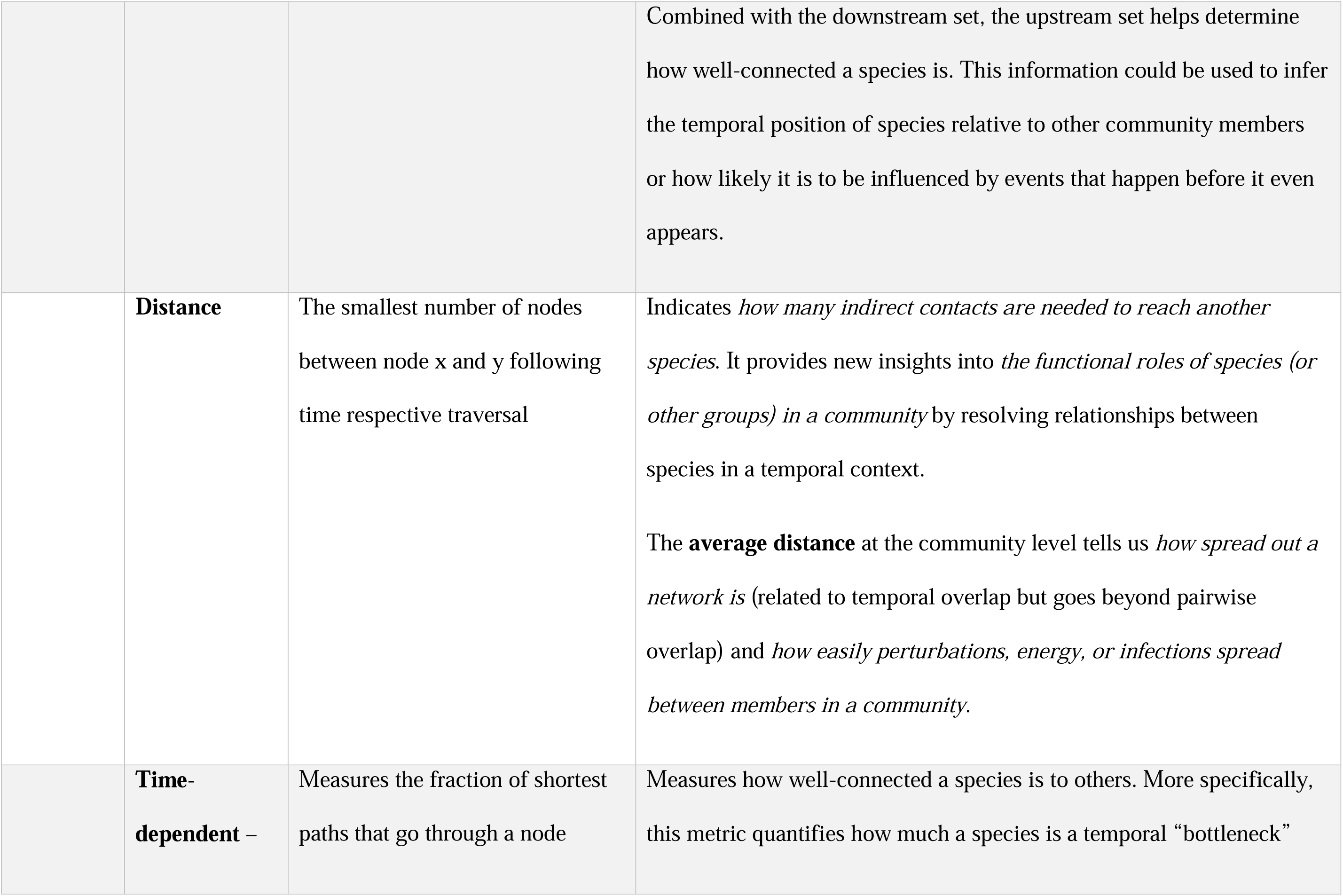
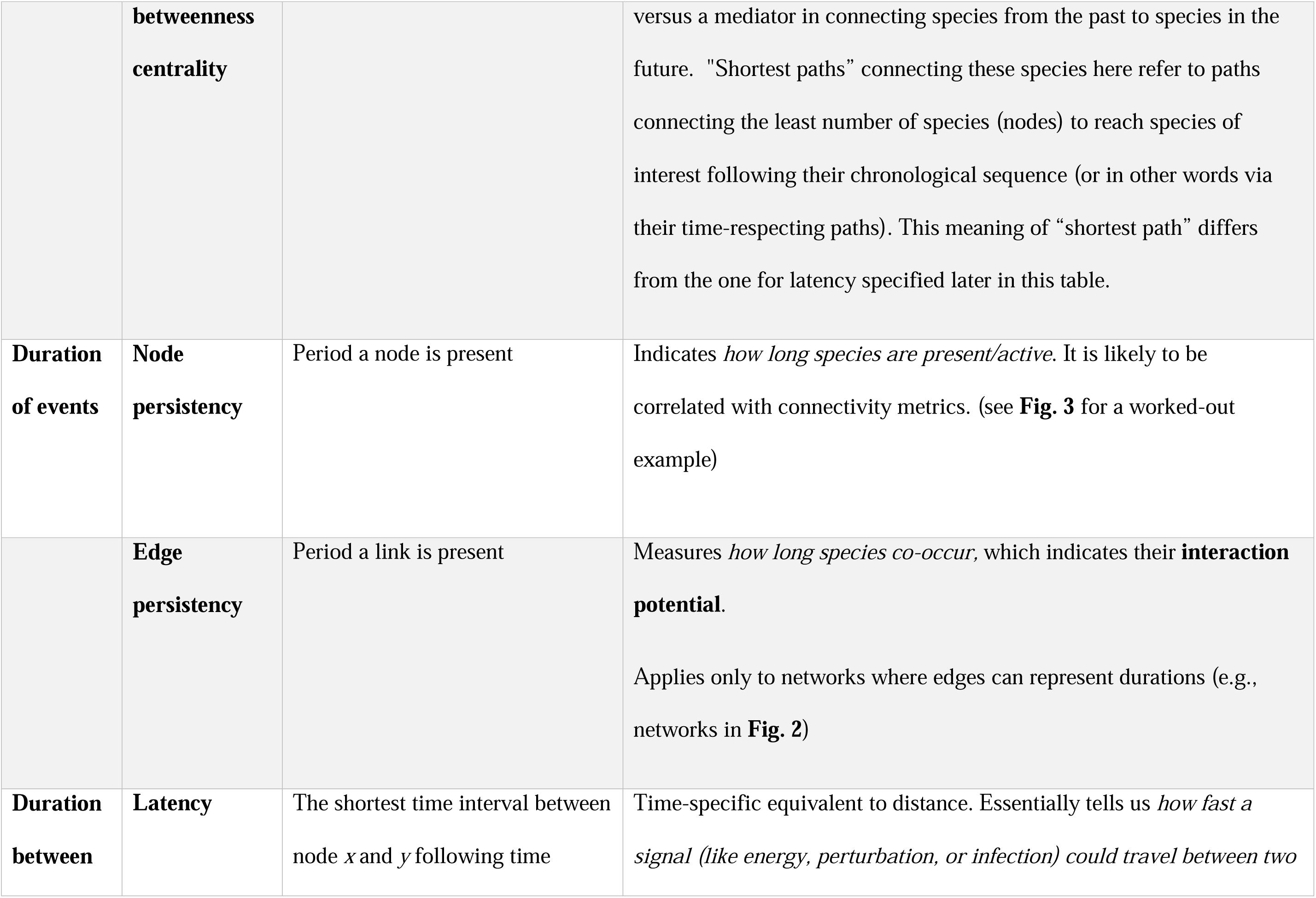
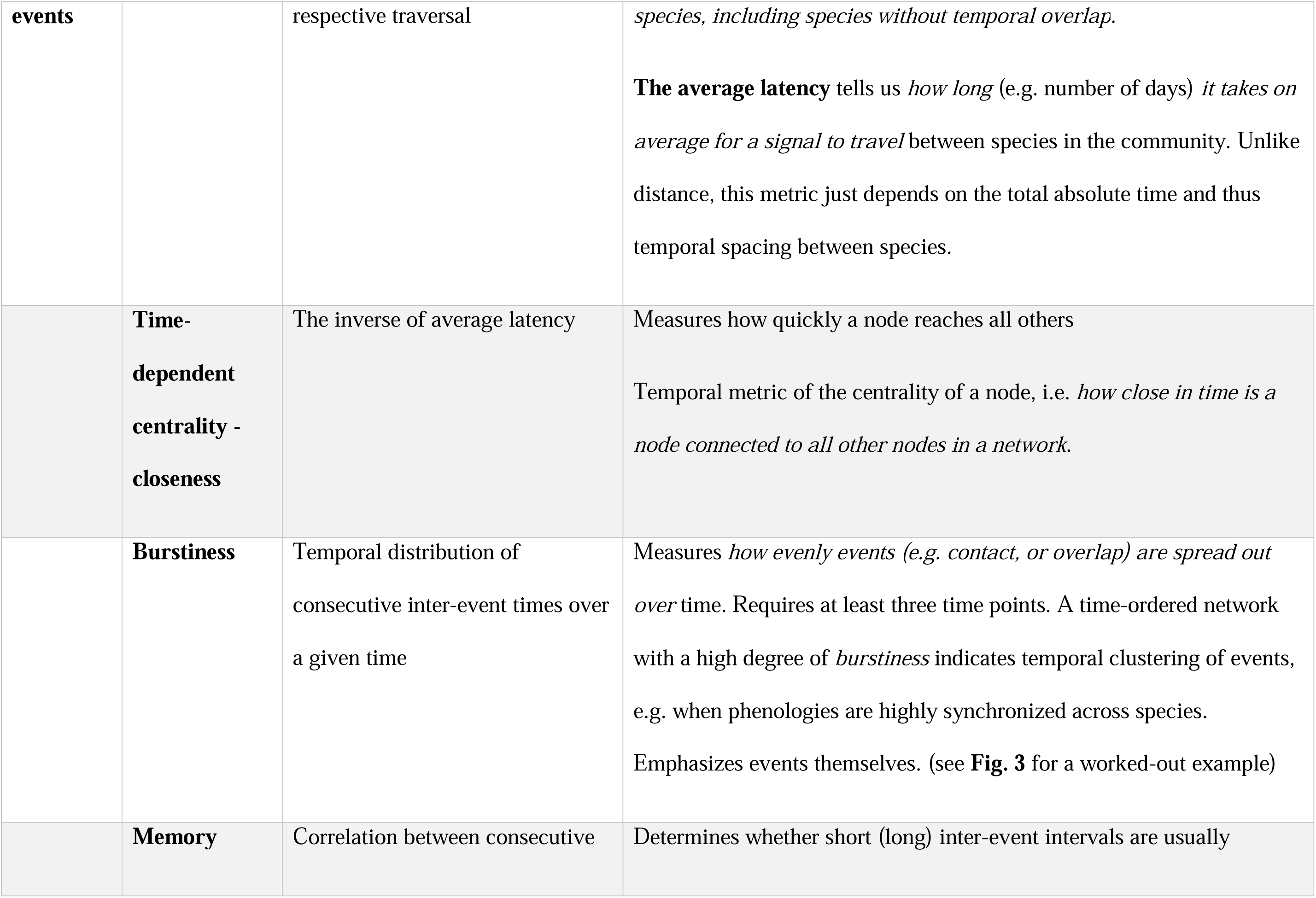
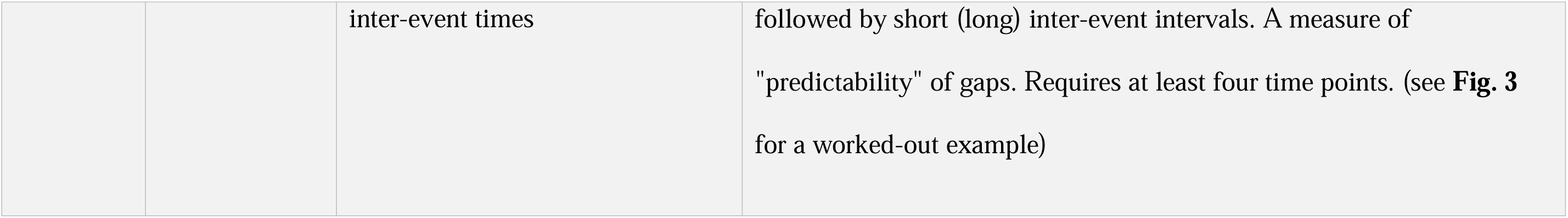
Metrics to quantify different aspects of the temporal structure of natural communities. For consistency, the meaning and applications column assumes the structure is a time-ordered species-co-occurrence (e.g., phenology) network. Although definitions vary among studies, these are all previously developed metrics and can be calculated with current software packages (see main text for details). Here, we largely follow the notations from (Holme & Saramäki 2019).

On the other hand, studies on temporal succession focus on turnover in species composition (e.g. Connell & Slatyer 1977; Dornelas *et al*. 2014; Van Allen *et al*. 2017; Penny *et al*. 2023; Poorter *et al*. 2023) and interactions (e.g. Díaz-Castelazo *et al*. 2010; Kantsa *et al*. 2018; Ushio *et al*. 2018; CaraDonna *et al*. 2021; Suzuki *et al*. 2023) and thus prioritize a multi-species perspective to capture how communities change over time. These studies revealed that species compositions and interactions are highly dynamic and can rapidly turn over across years, seasons, weeks, and even days, while some structural patterns at the network level (e.g., nestedness or connectivity) may change comparatively little (Petanidou *et al*. 2008; Kaartinen & Roslin 2012; Bramon Mora *et al*. 2020; Schwarz *et al*. 2020). These studies also highlight how we can infer important ecological and evolutionary processes that drive community dynamics (e.g., the concept of "succession" or "community drift") from the temporal organization of species. However, they still mainly rely on static metrics (either based on snapshots or time-aggregated data) that, at best, only contain indirect information on the sequence and duration of events. For instance, metrics for temporal turnover in species composition indicate that species come and go (e.g. Dornelas *et al*. 2014; Penny *et al*. 2023), but these metrics don’t provide information on the duration and sequence of events, e.g., how long species are present, which ones might return, how the current composition relates to previous events and if community states differed in how long they last. To use the music analogy again, temporal turnover, for example, helps quantify the tempo of a song but not the melody itself. Consequently, turnover can only capture a subset of patterns of the temporal structure itself. Of course, it would be unrealistic to assume that any single metric can comprehensively describe the complex temporal organization of communities. Accomplishing this requires a comprehensive toolset: a set of temporally explicit metrics, with each metric specialized to capture a different aspect of the dynamic temporal organization of communities.

Thus, despite significant recent advances that helped develop a more temporally explicit view of community ecology, we still have much to learn from the temporal structure of communities. A general framework accounting for the pros and cons of existing approaches would bring together different research areas in community ecology for a more comprehensive understanding of temporal community structure. Especially in this age of unprecedented environmental change, doing so should help the field better understand communities and anticipate how they will respond.

## A general approach to studying the temporal structure of natural communities: time-ordered co-occurrence networks

Here, we propose that time-ordered co-occurrence networks provide a powerful unifying framework to study the temporal structure of natural communities. Over the past decades, spatial ecology has proven that we can learn a lot from studying the (co-) occurrence patterns of species (Araújo *et al*. 2011; Borthagaray *et al*. 2014; Freilich *et al*. 2018; Gao *et al*. 2022; Galiana *et al*. 2023). The proposed framework takes a similar comparative approach for analyzing the temporal organization or species co-occurrences, except that it accounts for the directionality of time and keeps track of the possible causal chains of events.

### Time ordered networks

Time-ordered networks (also called time-varying, temporal, or dynamic networks) differ from traditional static networks because they are inherently dynamic and provide a complete record of the sequence and duration of events (Holme & Saramäki 2012; Masuda & Lambiotte 2020). In these networks, each node (vertex) and link (edge) is only active during specific time points (shown explicitly in **Fig. 2**). The unidirectionality of time (time always moves forward) means that the relationships between nodes have to follow a "time-respecting walk" of adjacent nodes and links. Importantly, these time-respecting walks have to follow the natural temporal sequence of events and thus lay out the possible causal chains contributing to community dynamics. For instance, if the path between nodes *A* to *B* ends before the link between *A* to *C* becomes active (**Fig. 2** scenario (a)), then node *B* could potentially indirectly affect node *C* (via *A*), but nothing from *C* can propagate to *B*, since signals cannot travel back in time. Thus, analyzing the structure of temporally-ordered networks helps us to better visualize and quantify the organization of events in time.

The study of time-ordered networks is ongoing, but it is a rapidly growing interdisciplinary field (Holme & Saramäki 2019; Masuda & Lambiotte 2020). Time-ordered networks, in general, have been used to describe temporal patterns for a range of systems, from public transportation and neurons in the brain to social networks (Holme & Saramäki 2019), but these networks have received surprisingly little attention in community ecology. Yet, as we will show below, they are ideally suited to study the temporal organization of natural communities, and recent advances and development of analytical tools not only help overcome the limitations of previous approaches discussed above but also provide unique opportunities for new temporally explicit analyses.

### Building time-ordered co-occurrence networks from phenology data

We will focus here on phenology data to demonstrate how we can apply this approach to quantify the temporal organization of species occurrences (but see "Future directions" for how this can be applied to other types of data). We already have a rich body of phenology data for various animal and plant communities in marine, freshwater, and terrestrial systems that are typically publicly available (e.g., USA National Phenology Network, Pan European Phenology project, Environmental Data Initiative). These datasets often have a fine temporal resolution and span decades, and they are rapidly growing with help from researchers, community-driven science initiatives, and government-funded observation networks (e.g., NEON, GBIF). Even with the taxonomic resolution varying by group (e.g. most vertebrates are resolved at species levels while some invertebrate groups are often only resolved at the genus or family level), collecting this data is being increasingly facilitated by recent technological advances. For instance, deploying cameras and machine learning to identify flowers and even pollinators at very fine temporal scales (Besson *et al*. 2022) can significantly help reduce the workload needed to collect this type of data. Similarly, DNA approaches can quantify phenologies and even interactions at similar or even finer temporal scales than direct observation (e.g., meta-barcoding of airborne pollen DNA, pollinator DNA in flowers, or eDNA analysis in freshwater and marine systems) (Besson *et al*. 2022; Jensen *et al*. 2022; Jønsson *et al*. 2023). Importantly, phenology data provide essential information about the temporal structure of communities by documenting when species are present and active in a given community.

Building time-ordered networks from phenology data (phenology networks) is relatively intuitive and generally straightforward in practice (see **Fig. 2** for a worked example). Following traditional approaches, we can build a network where a species represents a node, and its phenology determines the node’s time stamps (interval). Depending on the system, a node may thus have a single timestamp (e.g., present/active for a single day), a time interval (e.g., present from day x-y), or multiple time stamps if there is a temporal gap (pause) between activity periods (e.g., multiple distinct breeding events in a year). Links show relationships between nodes to represent when two species are active simultaneously in a given community. The time stamps for links would be similar to those of nodes (e.g., single or multiple time stamps), except that they represent the duration of the temporal overlap. In time-ordered networks, these links are sometimes called "contacts" to emphasize a time-limited information exchange (Holme 2015). Constructed in this way, these phenology networks capture co-occurrences and, thus, the complete temporal structure of communities.

Of course, if additional information on species interactions is available, it is straightforward to incorporate them into the network representation. Namely, we can restrict the topology of these co-occurrence networks further to focus only on direct interactions. Note that this would shift the focus from analyzing temporal co-occurrence patterns of species to analyzing temporal patterns of direct interactions. Furthermore, species or individuals could be grouped in other ways, e.g., by taxonomic group, traits, functional roles, or life stages. The taxonomic grouping may be necessary to standardize the data and facilitate interpretation when data are not fully resolved at the species level. Lastly, a particularly intriguing approach is to use this time-ordered network approach to track transitions in community states, with nodes representing community states and timestamped links signaling transitions between these states (see **Box 1 and Fig. 3** for an example). This approach is particularly promising in more complex systems and can help identify cycles, critical transitions in community states, and which species drive them. In general, the overall time-ordered network approach is very flexible, and the type of structure used to define nodes and links ultimately depends on the specific questions of interest and available information.

**Figure 3:**
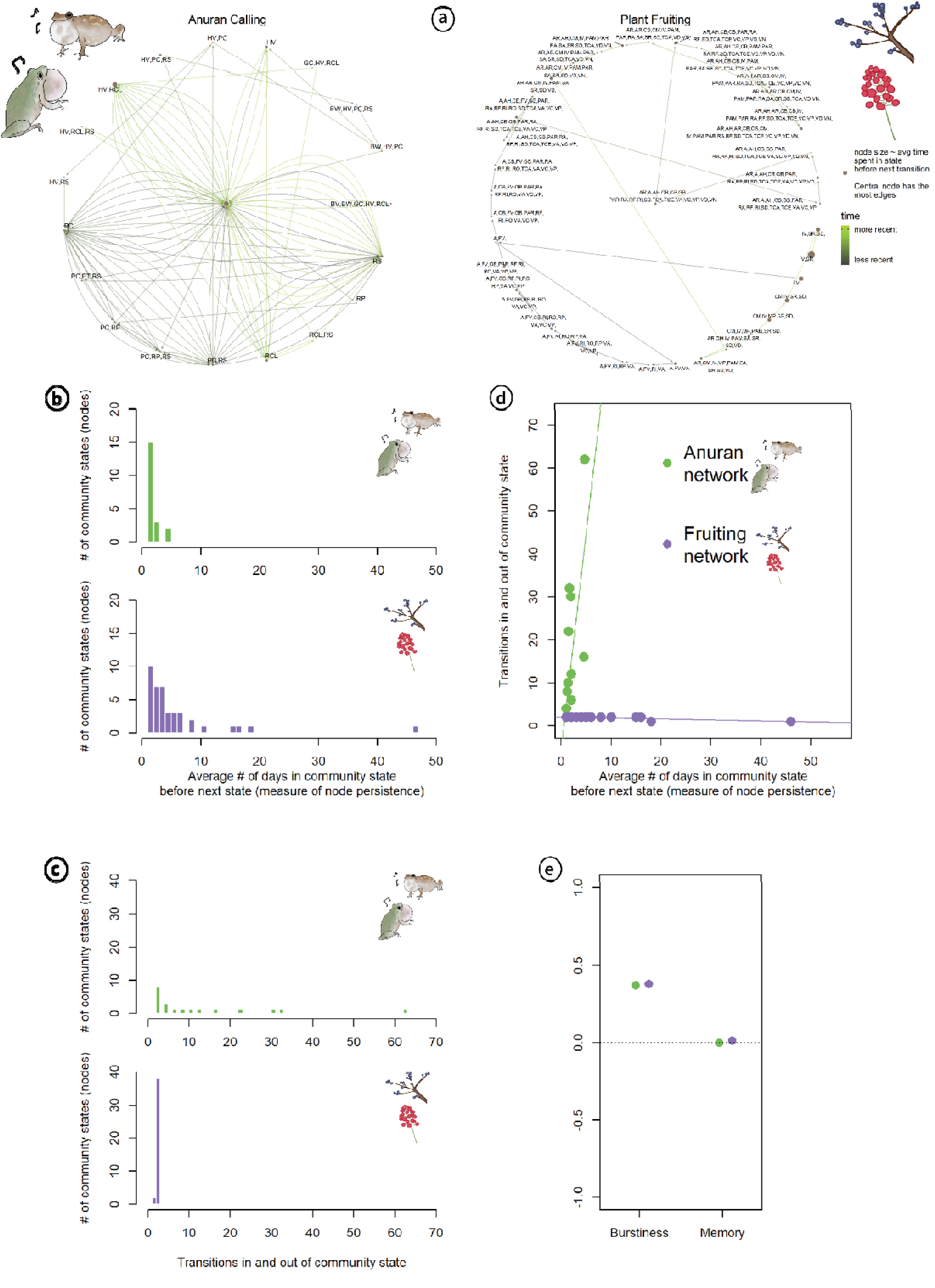
Changes in community structure over a 1-year time window in two different communities. **(a)** Transition networks, where nodes represent unique community states (species combinations, with each list of two-letter codes for a different species combination or an underscore for all species absent). The node’s size indicates its average duration (scaled to the respective community) and is scaled on a power function to improve visualization. Directed edges or links (arrows) indicate transitions between states. Note that we could have shown the same networks in a more condensed form by having a list of time stamps per arrow (like in Fig. 2). This would reduce the number of arrows drawn repeatedly between the same two community states, but we decided against this to visually emphasize the difference between the networks. The colors of links indicate the relative order of transitions (light green = near the end of the calendar year, dark brown = near the start of the calendar year). The anuran network (left) is based on daily calling data throughout an annual breeding season at one pond in Texas, USA, in the early 2000s (Carter *et al*. 2018), while the plant fruiting network (right) comes from Massachusetts, USA with single first and last date estimates based on observations make during 1850-1860 (Ellwood *et al*. 2022). Frequency plots show the distribution of **(b)** the duration (node persistency) of community states and **(c)** the number of transitions in and out of a given community state in the respective network (note that incoming and outgoing links count as two separate transitions), and **(d)** indicates how both metrics are correlated with each other within a network. **(e)** show two metrics that quantify different aspect of temporal distribution of events at the community level for each network: burstiness and memory (see **Table 2** for metric details).

### Quantifying time-ordered networks to understand natural communities

Converting species’ phenologies into time-ordered networks opens the door to calculating a range of informative metrics that could provide new and more profound insights into species’ biology, their temporal relationships, and the organization of natural communities. Some of these metrics for time-ordered networks are unique to temporal analyses, while others build on static network metrics to account for the unidirectionality of time. Here, we highlight and discuss a subset of previously developed time-ordered network metrics (**Table 2**) that we believe are among the most conceptually straightforward and biologically meaningful for analyzing phenology network data. Existing software can already calculate most of these metrics (e.g., tsna package (Bender-deMoll & Morris 2021), timeordered (Blonder 2015), and networkDynamic (Butts *et al*. 2023) in R, and Straph in Python). Several metrics like burstiness and memory have applications beyond network analysis and do not necessarily rely on network-specific software. Altogether, these metrics for phenological data capture important network properties such as connectivity, transmissibility, and persistence and the sequence and duration of events, the key features of temporal structure.

To better understand these metrics’ biological meaning and utility, it helps to think about what information they capture. Metrics like "*upstream set"* and "*downstream set"* (**Table 2**) (also called “backward reachable set” and “forward reachable set”) quantify how species are connected in time to other community members that appear either before or after them in time.

For instance, we can compare the *downstream set* across species to identify which species have the highest potential to influence other members within a given community or how infections might spread (see **Fig. 2** for worked-out example). Note, that these metrics depend on the community context and temporal position of species in the community; species that appear earlier will always have smaller upstream sets since few species appear before them (e.g. upstream set for species B versus species D in Fig. 2 panel a), and if there are temporal gaps in overlap, these sets will be interrupted and thus smaller (e.g. upstream set for species E panel a versus b).

Connectivity also depends on transmissibility—how easily and fast a signal or substance can travel across a community. In static networks, we typically measure this with the "*shortest path*", i.e., the smallest number of nodes we must traverse to connect one node to another. In time-ordered networks, this is sometimes called "*flow*" (Blonder *et al*. 2012), but distances between nodes can be measured in two ways in time-ordered networks: the number of nodes that need to be traversed (*distance*) or the actual time required to travel from one node to another (*latency*).

Comparing these metrics across species gives us a better idea of species’ relative temporal position in a given community. They could also help identify the potential for indirect effects between species (**Fig. 1**), differences in the sensitivity of communities to environmental perturbations (e.g., how fast perturbations or infectious diseases could spread in communities and how many species may be affected), or at what point in time communities are most vulnerable to perturbations. For instance, in communities with long average *distance,* perturbations will take a much longer time to reach all species and will have to go through a long temporal sequence of species which could dampen the effect of perturbation over time.

Similarly, indirect interactions are less likely between community members that are separated by long distances. Note, that these connectance metrics depend on the sequence and duration of and between events. Thus, they can give important insights into the functional role of species and their relationships in time (**Table 2**).

Other metrics, such as *burstiness, memory, node,* and *link* persistency (**Table 2**), focus on the duration of or between events to quantify the permanence/transience of time-ordered network structures, including temporal turnover of interactions. A high level of burstiness indicates temporal “clustering” (i.e. species tend to appear in groups during certain periods) and can help identify windows of opportunity (e.g., favorable environmental conditions, resource pulses) or temporal guilds. In contrast, a low level of *burstiness* indicates that the appearances (or presences) of species are more evenly spaced out over time and could indicate temporal niche partitioning (**Table 1**). Species with higher link persistency encounter more species over time and thus could indicate their interaction potential (Castro Arellano *et al*. 2010; Resasco *et al*. 2021), while node persistency could help identify different life history strategies among species (e.g., seasonal specialists with short node persistency versus generalists with long persistency).

We want to emphasize that time-ordered networks have not been widely applied to phenological data, so there is much room to grow and develop new metrics and tools. The metrics presented here are just a starting point, and we hope they inspire the development of complementary approaches for analyzing temporal communities. For instance, how can time-ordered networks incorporate count data from phenological datasets as weights to better predict ecological consequences of co-occurrences? How can we implement sequence in metrics of overlap or account for differences in proportional overlap? Developing these and other metrics is a promising new research area that will grow as we learn about temporal structure.

### Practical considerations

The size of any time-ordered network scales with the length of the period analyzed. Results of several temporally explicit metrics can differ based on the length of time over which network structures get quantified. This difference is particularly important for metrics that rely on time-respecting path lengths since they typically correlate positively with the period length (e.g., upstream and downstream set, distance, betweenness centrality in **Table 2**). Therefore, as best practice, any analyses of temporal community structure should have a clearly defined time window *t.* Fortunately, when analyzing phenology data, the calendar year or seasonal constraints (e.g., start and end of snow cover) often set the temporal window. However, these boundaries are sometimes blurred for some systems, e.g., when individuals or life stages stay active all year round. In this case, researchers must identify other biologically meaningful time windows (e.g., Does the system show a cyclical pattern reflecting the timing of El Nino Southern Oscillation events?) or stick with widely used time windows like the calendar year. In both cases, researchers must clearly communicate the time windows used in their analyses and standardize network metrics by the length of time windows when comparing time-ordered networks with variable time windows.

Visual representation of time-ordered networks can become cumbersome when exceeding a certain complexity. However, for metrics or network visualizations like those in **Fig. 3**, we can adapt existing core, open-source software packages developed for static networks (R packages: ggraph, igraph, network; Python packages: NetworkX). Given the increasing popularity of time-ordered networks and the development of new comprehensive guides (Masuda & Lambiotte 2020), we anticipate that more user-friendly options that work more directly with time-ordered networks will emerge over the next decade.

Finally, we must consider potential data limitations and caveats to help decide when this approach will be most helpful. Like all network approaches, it becomes more powerful when more nodes (species) are included. Similarly, very short or sparse time series may not contain enough information to reveal specific temporal patterns. Some metrics, like memory, require a minimum number of time points (**Table 2**). While phenology data is becoming increasingly available, the data recorded varies across studies, which determines the types of possible analyses. For instance, phenology datasets may only have start dates once per species per year (e.g., first flowering, first emergence). Without a biological notion of start and end times, a time-ordered network reduces to a temporal path or chronological sequence of events. This limits the amount of information we can extract from the temporal pattern. However, we can still extract some helpful information on temporal structure that can help put phenological shifts in a community context (see **Supplement, Fig. S1, S2)** for an example of an empirical application using a dataset on the leafout of woody plants).

## Applications and future directions

The acceleration of climate change has created a need for a more temporally explicit approach to ecology in the Anthropocene to understand and predict how these changes will affect natural ecosystems (Yang & Rudolf 2010; Wolkovich *et al*. 2014; Yang 2020). To predict how systems will change, we first need to understand the temporal structure of communities and the mechanisms that drive this structure. To date, progress in answering these questions has been hampered by a lack of a unified framework and limited available methods.

Our framework aims explicitly to overcome some of these limitations by offering comprehensive quantitative approaches that could scale across communities. With multiple metrics in hand, researchers can visually and quantitatively assess and compare the temporal structures of communities across timescales, space, and ecosystems (see **Box 1** and **Fig. 3** for a worked example) and help identify the underlying mechanisms. We have already mentioned some applications above but would like to highlight some additional exciting new research avenues and applications (**Box 2**). Of course, these areas serve as starting points and do not represent a complete list of potential applications.

### Functional roles and relationships among species

One novel aspect that emerges from this framework is that it can provide new ways to quantify and compare species’ "temporal niche" and functional roles by putting them in a temporally explicit community context. This analysis can reveal previously hidden functional differences between species within and across communities. For example, species that co-occur with the same number of species but differ in how long they are present in the community, when they are present, or how their interactions are distributed over time could show different responses to species extinctions and other environmental perturbations. To better understand species’ roles at the community level, we can perform robustness and sensitivity analyses (e.g., randomly removing species or species at risk of extinction) (Dunne & Williams 2009; Rudolf & Lafferty 2011) on these time-ordered networks to identify keystone species (e.g. those that result in most significant changes in time-ordered network metrics), or those species and ecosystems at greatest risk (i.e. species that experience the largest or more frequent changes in node-specific network metrics). These findings could complement existing knowledge to help guide management and conservation efforts in a rapidly changing world.

### General patterns and environmental drivers of temporal community structure

Since temporal structure is non-random and reflects a community’s ecological and evolutionary processes, we can expect to learn much from studying temporal patterns. For instance, perhaps weak burstiness and memory of fruiting plant communities (**Fig. 3e**) could arise from plant species starting but not ending their fruiting phenophases in the same month. Such a pattern suggests a potential shared underlying driver that might be linked to changes in environmental conditions or species’ life histories. However, weak burstiness could also arise from a temporal niche partitioning, e.g., to avoid competition for frugivores. These examples highlight how our framework can help identify new patterns and potential mechanisms and guide future research directions.

When we applied our approach to compare two very different communities (anuran calling versus a plant fruiting community), we found striking differences in the temporal network structure and some surprising similarities. These differences suggest corresponding patterns in the mechanisms regulating the temporal assembly of communities (see **Box 1** and **Fig. 3** for a complete example). This comparison highlights the power of having a set of metrics at hand to detect generalities and essential differences in the temporal structure across communities and environments that do not necessarily share the same abiotic drivers. At the same time, our worked examples also highlight significant gaps in our current knowledge of temporal community structure: ecology currently lacks a general framework to predict what these structures should look like or why and how they should vary across systems. Are those differences representative of the respective ecosystems or different environments? Do the similarities indicate shared drivers and biological constraints, or do they arise by chance? How will the patterns for a given type of ecosystem vary across different environments or with different species combinations? What will other types of communities look like? Answering these and many other related questions will significantly advance our understanding of the ecological and evolutionary processes shaping natural communities.

A crucial step toward answering these questions can involve comparing network metrics (**Table 2**) across systems, time, and space to identify potential drivers that shape the temporal structure of communities. For instance, our approach allows us to compare network metrics across latitudinal or altitudinal gradients to test how differences in seasonality influence the temporal organization of communities. Growing and reproductive seasons are often shorter in colder climates, but we don’t know how consistently this abiotic constraint shapes the temporal structure of communities. Do communities with shorter breeding/growing seasons show higher temporal overlap but shorter co-occurrences or the opposite? Does a shorter season alter the connectivity of time-ordered networks and their temporal relationships (e.g., *upstream set* versus *downstream set*, **Table 2**)? Are changes uniform across species, or do they depend on their temporal position (e.g., early versus late in the season)?

We can also see great potential in using our framework with temporally explicit null modeling approaches to narrow down potential mechanisms driving temporal structure. One of the challenges of working with any observational data, including temporal patterns, is that similar patterns can be created by different mechanisms. But, null models can often help us differentiate between mechanisms. For instance, we can randomize time stamps of nodes within a given community to break up sequence and temporal correlations or randomize the duration of phenophases to decouple the sequence and duration of events. The high degree of flexibility offered by these null or random network models allows for ensembles of increasingly randomized time-ordered networks useful for disentangling and identifying nested temporal patterns (Holme 2015). Comparing metrics from reference null models to empirical patterns within or across ecosystems helps identify non-random patterns in different aspects of temporal structure and better understand the processes that control temporal network structure. Say the latency between a set of species is much shorter than expected by chance. Then, there are processes causing temporal correlations between these species (Blonder *et al*. 2012). Ultimately, just like with spatial patterns (Gotelli & Graves 1996; Gotelli *et al*. 2010), we believe that confronting different null models with empirical data of time-ordered networks offers a particularly powerful approach for detecting general patterns in temporal community structure and for testing hypothesis to identify the underlying mechanisms of community assembly.

### Quantifying phenological shifts in a community context

Time-ordered networks also provide new tools to study the ecological consequences of phenological shifts. We want to reiterate that while past studies have primarily focused on metrics at the species level (i.e. first or mean arrival) or pairs of species (e.g. differences in first arrival), some effects may only become apparent in a community context. Small changes in the phenology of a species may have significant consequences at the community level depending on the temporal position of the species relative to other community members. For instance, the advancing phenology of species A in **Fig. 2** fundamentally alters the network structure and thereby affects the metrics (upstream and downstream set) of all species. In contrast, a similar shift in species D would have little consequence. Our approach represents a step toward identifying these species (e.g., species’ phenologies most important for maintaining the temporal structure and functioning of the existing ecosystem) and the groups likely to be the most affected by phenological shifts. These predicted differences in relative consequences following phenological shifts could then be used as *apriori* hypotheses and tested empirically.

### Paleo and bacterial data

Finally, we can also integrate time-ordered networks with other data types. For instance, recent advances in genetic methods have improved Quaternary paleo-records (Fordham *et al*. 2020), which have emerged as an important way to explore the long-term dynamics of communities and how they are linked to climate change (Blois *et al*. 2010; Jackson & Blois 2015). Our approach could easily be used with this type of data by simply using the presence of species in the archaeological record instead of phenologies to build time-ordered networks that study how communities have changed in the distant past. Since both types of data document co-occurrences, time-ordered networks could help bridge gaps between community ecology of the present and distant past, thereby improving our ability to understand the potential consequences of climate change (Fordham *et al*. 2020). Similarly, next-gen sequencing is increasingly providing us data on the temporal changes in the composition of bacterial communities (Espinoza *et al*. 2020). Combining this data with our time-ordered network approach could yield valuable new insights into the temporal structure of these communities, like their seasonal and annual dynamics, and how it is linked to their functioning (Deutschmann *et al*. 2023). Both paleo-records and sequencing data raise the importance of considering data types beyond those we have emphasized in the bulk of this paper to further strengthen our understanding of temporal community structure.

## Conclusions

Like spatial ecology, integrating temporal structure into community ecology has the potential to fundamentally alter our understanding of the critical forces and processes that shape natural communities. To date, we still lack a general understanding of how temporal structures of communities vary, what general patterns might exist in nature, and what the underlying drivers are. We know even less about their consequences. Nevertheless, answering these questions is fundamental to understanding the maintenance of ecological communities (Ushio et al. 2018) and predicting how they will respond to future perturbations and climate change. Our framework outlines how to take the next important steps toward building a temporally explicit framework for community ecology and suggests new and exciting opportunities to help overcome some previous limitations. Building this framework is particularly important to understand and predict how natural ecosystems will change as climate change reshuffles the temporal structure of natural communities worldwide. The next challenges include collecting, accessing, and analyzing this temporal data to shed new light on this understudied dimension of natural systems.

## Acknowledgments

We would like to thank A.E. Dunham, J. HilleRisLambers, A. Silva, and A. Simha for feedback on the ideas, methods, and manuscript. This work was supported by NSF DEB-1655626 to V.H.W. Rudolf and by the NSF Graduate Research Fellowship Program under Grant No. 1842494. Any opinions, findings, and conclusions or recommendations expressed in this material are those of the author(s) and do not necessarily reflect the views of the National Science Foundation.

### Box 1: Applying the temporal framework: temporal patterns of anuran calling community versus plant fruiting community

To highlight some of the utility of our framework, we show an example of how we can use it to compare the temporal structure of communities using publicly available datasets on two very different systems: one on anuran calling in Texas (Carter *et al*. 2018) and the other on fruiting phenology in Thoreau’s Concord (Ellwood *et al*. 2022) (**Fig. 3**). Due to the complexity of the communities and to highlight the flexibility of our approach, we focused here on changes in community states. The approach to building time-ordered networks is still the same (see **Fig. 2**), except that here, nodes represent a unique combination of co-occurring species (community state), and links (relationships between nodes) show the transition between the different community states. These links have time stamps for when transitions occur, while nodes have time intervals for the duration of community states.

We can immediately see striking differences in the temporal structure of both networks (**Fig. 3a**). The anuran-calling time-ordered network shows many recurring community states but with considerable variation in how long each state lasts and how often it reappears (green histograms in **Fig. 3b** and c, respectively). This pattern arises because most species have more than a single time interval during which they are active and can vary in the length of their phenophases (i.e., few species with long vs many with short phenophases). Together, this creates a complex and highly dynamic network with many possible causal chains and where species with long phenophases (which form the long tail in the histogram in **Fig. 3b**) act as "hubs" connecting different community states. In contrast, the fruiting network suggests a sequential turnover of community states because species typically only fruit during a single period. Consequently, there is a much more explicit temporal hierarchy with a simpler causal structure (who comes first and could influence later arrivers). Each community state typically occurs only once (note that because in and out transitions count each separately, this pattern is indicated by the step peak at 2 in **Fig. 3b**). In both networks, most states (nodes) do not last long before transitioning to another state, but a minority of states in the plant fruiting network last much longer and form a right-skewed distribution. When plotting the distributions of how long each community state lasts on average (node persistence) and recurring community states in the same plot (**Fig. 3d**), we see the networks follow very different relationships reflective of differences in time-ordered network structure (i.e., strongly positive relationship in frog network versus weak to no relationship in plant network). Yet, despite these differences, both communities show similar temporal distribution of events at the community level, i.e., a moderate signal of temporal clustering of community shifts (burstiness in **Fig. 3e**) and no clear autocorrelation in times between community shifts (memory in **Fig. 3e**). This moderate clustering suggests some periods when multiple species tend to enter/leave in quick succession (e.g., due windows of opportunities or species belonging to similar functional guilds). These examples illustrate just some of the various analyses that can be performed with our approach to comprehend better the common and unique temporal structures and functions of natural communities.

### Box 2: Some of the many outstanding questions (in no particular order) that can be addressed with the framework proposed here (see “Applications and Future directions” for worked examples)

- What are the general, non-random features of temporal community structure that are shared across most ecosystems?
- How do environmental conditions and patterns in human land use influence different aspects of the temporal structure of communities?
- How do ecological and evolutionary processes shape the temporal structure of communities?
- What effects do seasonal constraints (like the length of the growing season) have on the temporal structure of communities?
- How does the temporal structure of communities change across time and space?
- How will climate change (including phenological shifts) alter the temporal structure of communities?
- How are species’ traits linked to the temporal structure of communities?
- Which causal relationships or pathways of flow (or spread) are plausible in a community?
- How does species, functional, or trophic diversity influence the temporal structure of communities?
- How is the temporal structure of communities linked to their stability?
- What new metrics and tools are missing to better capture the temporal structure of communities?

